# Neuromodulator-induced Temperature Robustness in a Motor Pattern: a Comparative Study Between two Decapod Crustaceans

**DOI:** 10.1101/2023.11.23.568465

**Authors:** Wolfgang Stein, Carola Städele

## Abstract

While temperature fluctuations pose significant challenges to the nervous system, many vital neuronal systems in poikilothermic animals function over a broad temperature range. Using the gastric mill pattern generator in the Jonah crab, we previously demonstrated that temperature-induced increases in leak conductance disrupt neuronal function and that neuropeptide modulation provides thermal protection. Here, we show that neuropeptide modulation also increases temperature robustness in Dungeness and Green crabs. Like in Jonah crabs, higher temperatures increased leak conductance in both species’ pattern-generating neuron LG and terminated rhythmic gastric mill activity. Likewise, increasing descending modulatory projection neuron activity or neuropeptide transmitter application rescued rhythms at elevated temperatures. However, decreasing input resistance using dynamic clamp only restored the rhythm in half of the experiments.

Thus, neuropeptide modulation increased temperature robustness in both species, demonstrating that neuropeptide-mediated temperature compensation is not limited to one species, although the underlying cellular compensation mechanisms may be distinct.

**Summary statement:** This study shows that the release of neuropeptide from modulatory projection neurons plays a crucial role in maintaining neuron and circuit function at elevated temperatures across crustacean species.

## Introduction

Temperature changes are a particular challenge for the nervous system, as they can disrupt the well-balanced weighting of intrinsic and synaptic ionic conductances that are required to maintain neuronal function and behavioral performance (Robertson and Money, 2012, Harding et al., 2019, Marder and Rue, 2021, Robertson et al., 2023). Poikilothermic aquatic animals are especially prone to temperature changes because they do not actively maintain their body temperature, live in an environment with high heat transfer, and are especially susceptible to climate change-induced global warming and the resulting heatwaves (Reid et al., 2009, Somero, 2010, Somero, 2012, Mann et al., 2018, Oliver et al., 2018). Nevertheless, many vital behaviors in these animals function across a wide temperature range, making them ideal for studying the mechanisms of temperature resilience. Recent studies have revealed that some neural circuits exhibit intrinsic robustness to acute temperature perturbations arising from cellular and synaptic ionic conductances (Alonso and Marder, 2020). Other neuronal circuits rely less on intrinsic properties but gain robustness through external neuromodulatory influences. In particular, the gastric mill central pattern generator in the stomatogastric nervous system of the Jonah crab *Cancer borealis* (Stein, 2017) shows very little intrinsic robustness against heating. A change of only a few degrees Celsius can terminate the gastric mill motor pattern. We have previously shown that this termination is due to an increase in membrane leak conductance with temperature in the lateral gastric (LG) neuron, which eliminates spike activity and shunts LG bursting (Städele et al., 2015, DeMaegd and Stein, 2021). Since LG is a member of the gastric mill half-center circuit that drives the gastric mill rhythm, the rhythm terminates. However, we have also shown that neuropeptide modulation enables thermal robustness and allows normal neuronal functioning at much more elevated temperatures. Specifically, additional substance P-related peptide CabTRP Ia (*Cancer borealis* tachykinin-related peptide Ia) enables neurons in the gastric mill central pattern generator to produce bursts of action potentials at elevated temperatures in *C. borealis*. CabTRP Ia is released from a pair of descending projection neurons (the modulatory commissural neurons 1, MCN1), and there are indications that MCN1 activity increases with temperature (Städele et al., 2015, Städele and Stein, 2022). CabTRP Ia rescues neuronal activity by activating a voltage-gated inward current (I_MI_) that counterbalances temperature-induced increases in leak currents. Since neuromodulator release from descending projection neurons is a common process to control neuronal activity in many circuits and across taxa, we hypothesized that neuropeptide-mediated temperature robustness is not distinctive to *C. borealis*. This study aims to test this hypothesis by comparing the effects of temperature and peptide neuromodulation on LG neurons in Dungeness crabs (*Cancer magister*) and green crabs (*Carcinus maenas*) with findings from previous studies on *C. borealis*. Each species inhabits different environments and experiences different temperature fluctuations than the previously studied *C. borealis*.

## Materials and Methods

### Animals

Adult *Cancer magister* were purchased at a local store and kept in filtered, aerated artificial seawater at 11°C until used. Adult *Carcinus maenas* crabs were provided by Dr. Markus Frederich (University of New England, Maine, USA) and Adam Smith (Perfect Limit Green Crabs, Massachusetts, USA). *C. maenas* crabs were kept in filtered, aerated artificial seawater at 20°C until used. Figure 1A gives an overview of the species used, where they originated, and the temperatures they typically encounter in their habitat.

**Figure 1:**
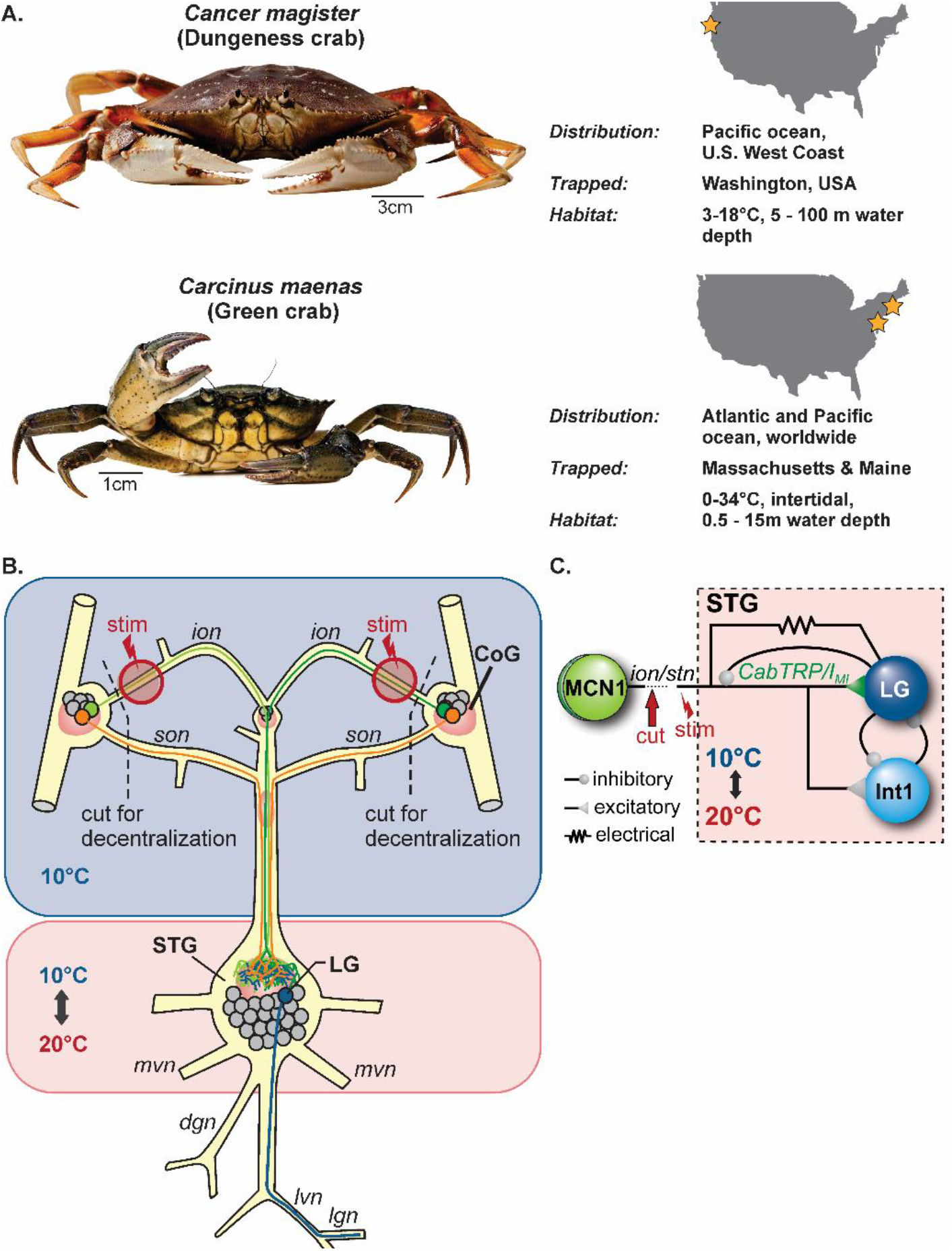
(**A**) The two crab species used in this study, their habitat, and their point of origin (Prentice and Schneider, 1979, Logan and Frederich, 2017). B. The temperature of the STG (containing the gastric mill pattern generating circuit) was changed by superfusing temperature-controlled saline (10 to 20°C). CabTRP Ia (0.1 -1 μM) was superfused exclusively to the STG. Upstream areas, including descending projection neurons in the paired commissural ganglia (some indicated in color), were kept in regular saline at constant temperature (∼10°C). Dashed lines indicate nerves that were cut for decentralization. (**B**) We recorded intracellularly from LG, which controls the main protractor muscles in the gastric mill. In some experiments, we extracellularly stimulated axons of descending projection neurons within the paired inferior oesophageal nerves (stim). (**C**) The circuit activating LG has been identified in *Cancer borealis*: the descending modulatory projection neuron MCN1 excites and modulates LG activity via an electrical synapse and by paracrine release of the peptide CabTRP Ia. Abbreviations: STG: stomatogastric ganglion, CoG: commissural ganglion, LG: lateral gastric neuron, Int1: Interneuron 1, *ion*: inferior oesophageal nerve, *son*: superior oesophageal nerve, *mvn*: medial ventricular nerve, *dgn*: dorsal gastric nerve, *lvn:* lateral ventricular nerve, *lgn:* lateral gastric nerve, *stn*: stomatogastric nerve.

### Dissection

Established protocols were used. After ice anesthesia for 20 - 40 minutes, the stomatogastric nervous system was isolated according to (Gutierrez and Grashow, 2009), pinned out in a silicone-lined (Wacker) petri dish, and continuously superfused with physiological saline (10°C). We worked with fully intact and decentralized STNS preparations. In the latter, the stomatogastric ganglion (STG) was separated from the commissural ganglia (CoGs) by transecting the paired *ion*s and *son*s (Fig. 1B).

The animal species used in this study are not subject to ethics approval at Illinois State University. We nevertheless adhered to general animal welfare considerations regarding humane care and use of animals. Water quality, salinity, and temperature were monitored daily. Crabs were sacrificed using ice, which is a method recognized as acceptable under the AVMA guidelines for the euthanasia of aquatic invertebrates.

### Solutions

*C. magister* saline was composed of (in mM) 440 NaCl, 26 MgCl_2_, 13 CaCl_2_ x 2H_2_O, 11 KCl, 10 Trisma base, 5 maleic acid, pH 7.4-7.6 (Sigma Aldrich). *C. maenas* saline was composed of 410 NaCl, 11 KCl, 11 CaCl_2_ x 2H_2_O, 21 MgCl_2_ x 6H_2_O, 30 NaHCO_3_, 11 Trizma base, 5.1 Maleic acid, pH 7.4-7.6 (Sigma Aldrich). CabTRP Ia (GenScript) was added to the saline in some experiments. Solutions were prepared from concentrated stock solutions immediately before the experiment. Stock solutions were stored at -20°C in small quantities. Measurements were taken after 15 minutes wash in.

### Electrophysiology

Extracellular action potentials were recorded, filtered, and amplified with an AM Systems amplifier (Model 1700). We recorded from several gastric mill motor nerves, including the lateral gastric nerve *lgn*, which contains the LG axon, the dorsal gastric nerve *dgn*, which contains the DG axon, the medial ventricular nerve (*mvn*), and the lateral ventricular nerve *lvn* to monitor pyloric and gastric mill rhythm activity (Fig. 1B). Intracellular recordings were obtained from STG cell bodies using 20-30 MΩ glass microelectrodes (Sutter 1000 puller, 0.6 M K_2_SO_4_ + 20 mM KCl solution, or cytoplasm matched solution: 20 mM NaCl, 15 mM Na_2_SO_4_, 10 mM Hepes, 400 mM potassium gluconate, 10 mM MgCl_2_ (Hooper et al., 2015). Signals were filtered and amplified through an Axoclamp 900A amplifier (Molecular Devices) in bridge or two-electrode current clamp mode. Files were recorded, saved, and analyzed using Spike2 Software at 10 kHz (version 7.18; CED) and a Power 1401 (CED). Input resistance was measured using hyperpolarizing current pulses (-2nA, 2s duration). Membrane potential voltage deflections were measured in steady-state (after 2s).

To elicit gastric mill rhythms in decentralized nervous system preparations, we extracellularly stimulated the axons of descending projection neurons in the part of the transected *ion* that remained connected to the STG (marked with stim in Fig. 1B) at 10°C. Both *ions* were tonically stimulated for 200s with the same frequency (Master-8 stimulator (AMPI), 1 ms pulse duration). In *C. borealis*, the lowest activation threshold of the *ion* elicits action potentials in the MCN1 projection neuron (Coleman et al., 1995, Bartos and Nusbaum, 1997). Tonic MCN1 stimulation induces a specific version of the gastric mill rhythm that has been thoroughly characterized in previous studies (Coleman et al., 1995, Stein et al., 2007, Hedrich et al., 2011). In the two species studied in this paper, the presence of LG PSPs during *ion* stimulation was used to confirm that the activation threshold was reached. We adjusted the activation threshold separately for each *ion* by slowly increasing stimulation voltage until PSPs were obtained in LG. Once the activation threshold was reached, it was maintained at all temperatures. It is unclear whether the axonal projections in the two species studied here are identical to those in *C. borealis*. However, stimulating at subthreshold amplitudes did not elicit any response of the pyloric or gastric mill neurons, suggesting that, like in *C. borealis*, threshold stimulation recruited an MCN1-like descending projection that innervates LG. Stimulation frequency was increased in 1 Hz steps until a gastric mill rhythm was observed at 10°C (=*threshold frequency*).

### Temperature manipulations

Preparations were continuously superfused with physiological saline. Temperature was manipulated inside of a petroleum jelly well around the STG. The well thermally isolated the STG from the rest of the nervous system. The temperature inside and outside the well was controlled independently with two saline superfusion lines, cooled by separate Peltier devices. Temperature was continuously measured within a few millimeters of the STG and the CoGs with separate temperature probes (Voltcraft 300K). We selectively altered STG temperature while the surrounding nervous system was kept constantly at ∼10°C. Temperature was changed by ∼1°C/min unless otherwise mentioned. With intracellular recordings, the temperature was changed by ∼1°C/6-7 min to avoid losing the recording. Measurements were taken in steady-state, i.e., after keeping the temperature constant for at least two minutes.

### Dynamic clamp

(Sharp et al., 1993) was used to reduce or increase LG input resistance. This was achieved using Spike2 software and two-electrode current clamp mode. In each preparation, input resistance and resting potential were measured at cold temperature and at the elevated temperature at which the rhythm had crashed to determine temperature-induced changes. Changes in input resistance were implemented by an artificial leak conductance computed in dynamic clamp:

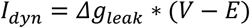

where *I*_*dyn*_ is the injected current and *Δleak* represents the difference in leak conductance between cold and elevated temperatures. *E* was taken as the resting potential at elevated temperature. *E* and *Δleak* were calculated separately for each preparation. Custom-written Spike2 scripts and sequencer files were used to implement the equation and control the 1401 AD board.

### Data analysis

Figures were prepared with CorelDraw X7 (Corel Cooperation), Excel 365 (Microsoft), and SigmaPlot (version 15, Jandel Scientific). Analyses were carried out using Spike2 (CED, UK). Unless stated otherwise, data are presented as mean±SD. Alternatively, individual data points for each animal are given. Significant differences are stated as *p<0.05, **p<0.01, ***p<0.001.

## Results

### Influence of temperature on the gastric mill rhythm

We tested the influence of temperature on the gastric mill rhythm by controlling the temperature of the saline superfusion to the STG. At the same time, upstream areas, including the paired commissural ganglia (CoGs), were kept at a constant temperature (∼10°C) (Fig. 1B). The gastric mill rhythm is a biphasic, episodic pattern, and in *C. borealis* is known to be driven by the half-center oscillations of Interneuron 1 (Int1) and the LG neuron (Fig. 1C) (Stein et al., 2007). LG and Int1’s cell bodies are in the STG, but rhythmic gastric mill activity requires modulatory input from descending projection neurons in the upstream commissural ganglia (CoGs, Figs. 1B, C). These projection neurons can be spontaneously active but are strongly activated by a diverse set of sensory pathways (Nusbaum and Beenhakker, 2002, Beenhakker and Nusbaum, 2004, Blitz et al., 2004, Blitz et al., 2008, Hedrich et al., 2009, Diehl et al., 2013, Nusbaum, 2013, Stein, 2017). In *C. borealis*, they can also be selectively stimulated to elicit specific versions of the gastric mill rhythm (Coleman et al., 1995).

Our previous data from *C. borealis* demonstrated that LG, without which the gastric mill rhythm cannot function, stops producing rhythmic bursts when the STG temperature is raised by only a few degrees Celsius(Städele et al., 2015, DeMaegd and Stein, 2021, Städele and Stein, 2022). In a first set of experiments in *C. magister*, we tested the effects of temperature on spontaneously occurring gastric mill rhythms. As previously reported in *C. borealis* (Städele et al., 2015, DeMaegd and Stein, 2021, Städele and Stein, 2022), these rhythms stopped when the STG was selectively heated (Fig. 2A).

**Figure 2:**
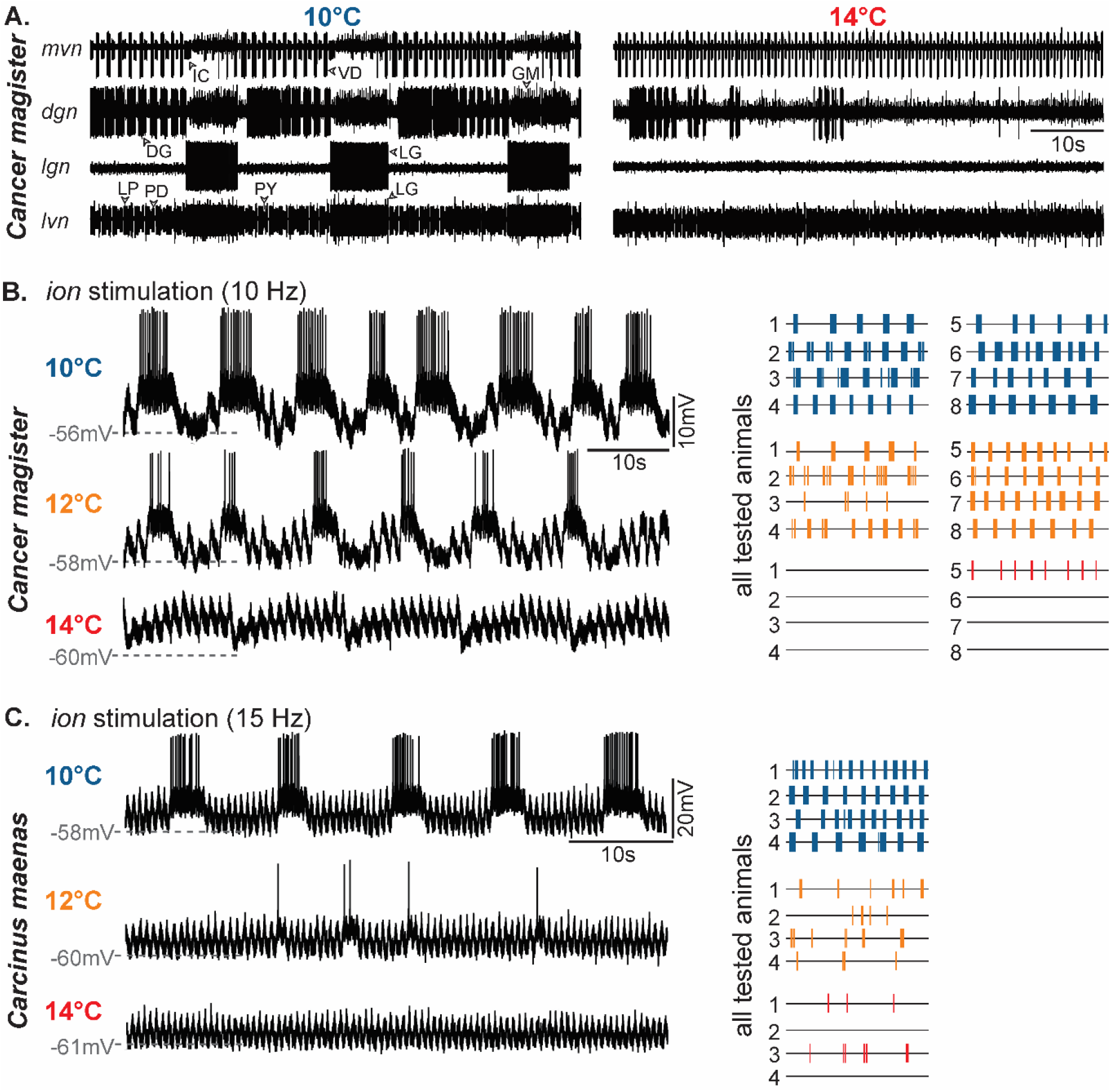
C. magister and C. maenas gastric mill rhythms terminate at elevated temperature. (**A**) Extracellular nerve recordings showing the spontaneous gastric mill rhythm at low (10°C) and elevated temperature (14°C). Four extracellular recordings are shown. Top: medial ventricular nerve (*mvn*) showing the activities of the inferior cardiac (IC, small unit) and ventricular dilator (VD, large unit) neurons. Second from top: dorsal gastric nerve *dgn* showing the activity of the dorsal gastric (DG; large unit) and gastric mill (GM, small units) neurons. DG is a functional antagonist of LG and the GM neurons. Third from top: lateral gastric nerve *lgn*, showing the activity of LG. Bottom: lateral ventricular nerve *lvn*, showing the pyloric rhythm. The pyloric rhythm is a fast triphasic rhythm continuously active and shows a much larger intrinsic temperature robustness than the gastric mill rhythm. At 14°C, LG activity ceases, and the gastric mill rhythm terminates while the pyloric rhythm continues. Recordings are from the same preparation. LP: lateral pyloric neuron, PD: pyloric dilator neurons, PY: pyloric constrictor neurons. (**B**) Intracellular recording of *C. magister* LG during continuous extracellular *ion* stimulation with 10 Hz at 10, 12, and 14°C. LG was rhythmically active at 10 and 12°C, but activity ceased at 14°C. Right: Representation of LG spike activity for all preparations tested at 10, 12, and 14°C. Each trace shows 100 s during continuous ion stimulation, with each vertical line representing an action potential in LG. (**C**) Intracellular recording of *C. maenas* LG during continuous extracellular *ion* stimulation with 15 Hz at 10, 12, and 14°C. LG was rhythmically active at 10°C, but activity ceased at 12 and 14°C. Right: Representation of LG spike activity for all preparations tested at 10, 12, and 14°C.

Spontaneous gastric mill rhythms can vary in activity patterns because multiple descending projection neurons may contribute to their initiation. To investigate the temperature effects on the rhythm better, we transected the axons of all projection neurons (see Material and Methods) and stimulated both *ions*. In *C. borealis*, this elicits a distinct and well-described version of the rhythm (Bartos et al., 1999, Hedrich et al., 2011) that is driven by the paracrine release of CabTRP Ia from MCN1 onto LG (Stein et al., 2007), in addition to providing excitatory synaptic input via an electrical synapse (Stein et al., 2022). We recorded intracellularly from LG in these experiments. Like in *C. borealis*, transecting the axons of the projection neurons terminated ongoing spontaneous gastric mill rhythms, and stimulating the *ion* consistently elicited rhythmic activity in LG when the STG was kept at 10°C (Fig. 2B). The elicited rhythm strongly resembled the MCN1-type gastric mill rhythm of *C. borealis*, as evidenced by the burst patterns and subthreshold membrane potential oscillations (Coleman et al., 1995). Increasing the temperature reversibly terminated rhythmic bursting in LG (Fig. 2B). This was consistently the case at 14°C.

In *C. maenas*, no spontaneous gastric mill rhythms were observed (N=19). However, stimulation of the *ion* at 10°C elicited a gastric mill rhythm (Fig. 2C), albeit less reliably than in *C. borealis* and *C. maenas*. In 8 of 12 experiments, no gastric mill rhythm could be elicited. Failures to elicit gastric mill rhythms occasionally also occurred in *C. magister*, but only in less than 10% of preparations. The inability to elicit a gastric mill rhythm in *C. maenas* was not due to a failure of the stimulation. Clear postsynaptic potentials from each stimulus pulse were seen in the LG neuron, even when no rhythm was elicited, indicating that the MCN1-like projection neuron was being activated. In addition, our stimulation had a clear excitatory effect on the pyloric rhythm, further suggesting that the stimulation was successful. One of the reasons for these failures to activate the gastric mill rhythm could have been that we were stimulating at the wrong temperature. We thus changed the STG baseline temperature to various temperatures between 10 and 20°C in preparations that did not show a gastric mill rhythm at 10°C. However, no gastric mill rhythm could be started in these preparations in the tested temperature range. Finally, increasing the stimulation frequency did not establish a gastric mill rhythm either.

In those preparations where a gastric mill rhythm could be elicited, the rhythm was stable and showed similar activity patterns as in *C. magister* (Fig. 2C) and as previously reported in *C. borealis* (Coleman et al., 1995). Increasing the STG temperature by 4°C ended burst generation in LG and stopped or disrupted the rhythm (N=4, Fig. 2C).

### Temperature effects on cellular properties of the LG neuron

To investigate the cellular effects of heating the STG on LG, we recorded intracellularly from the LG soma and measured the changes in resting membrane potential, synaptic input, and action potential amplitude. In *C. magister*, the resting membrane potential hyperpolarized when the STG temperature was increased (Fig. 3A). Concurrently, action potential amplitudes diminished with higher temperatures. Similarly, the amplitude of the postsynaptic potential (PSP) elicited by each stimulus pulse, i.e., by each MCN1-like action potential, diminished.

**Figure 3:**
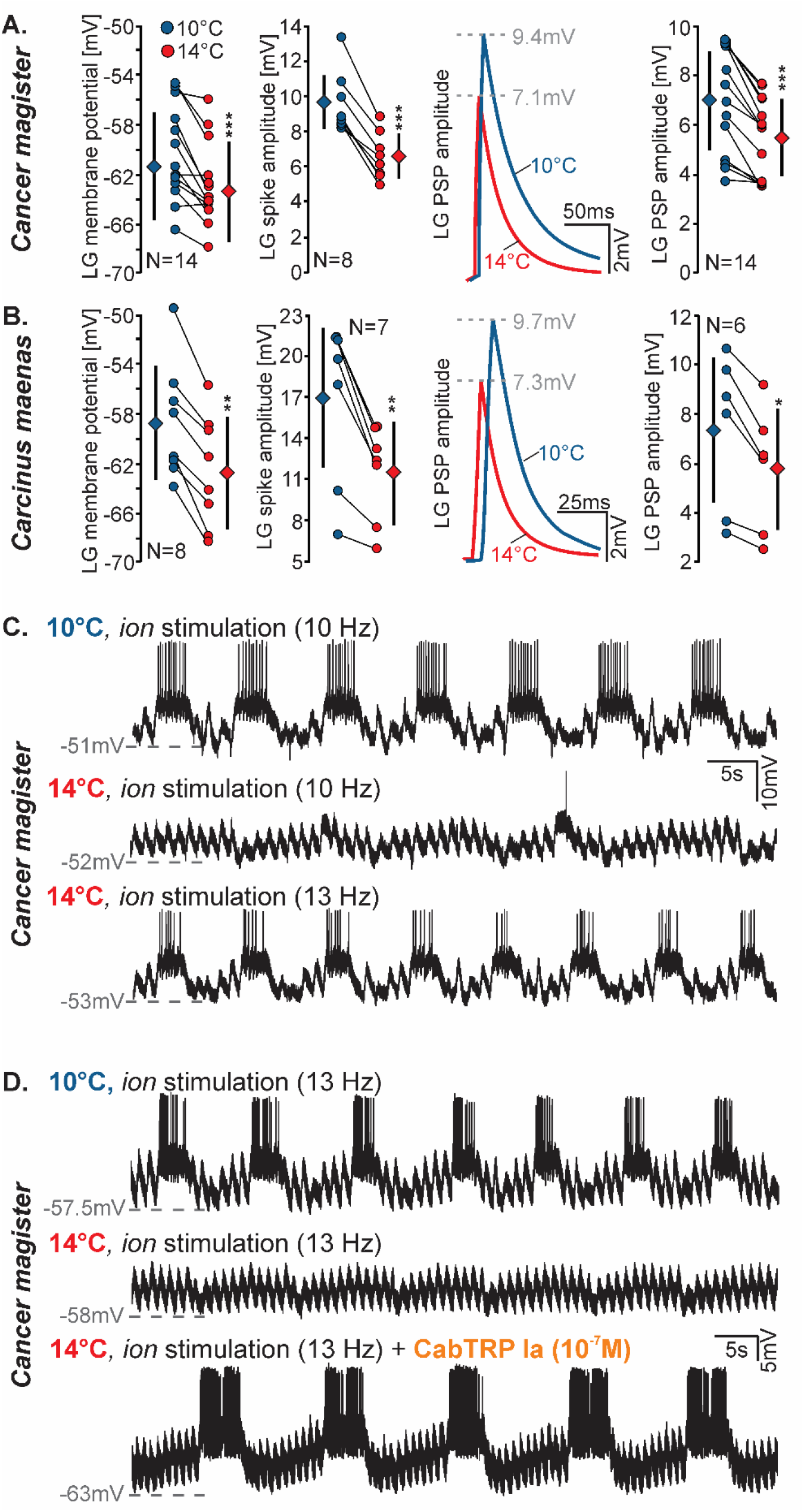
Comparison of temperature effects on LG’s cellular properties in *C. magister* (**A**) and *C. maenas* (**B**). *P<0.05; **P<0.01; ***P<0.001; paired t-test. Second from right: Comparison of original recordings of the MCN1-like elicited electrical PSPs in LG at 10 (blue) and 14°C (red). (**C**) Increasing *ion* stimulation frequency can rescue a crashed *C. magister* gastric mill rhythm at elevated temperature. Intracellular recording of LG during continuous extracellular ion stimulation with 10 Hz at 10 and 14°C. LG was rhythmically active at 10°C, but activity ceased at 14°C. Increasing *ion* stimulation frequency to 13 Hz re-established the rhythm at 14°C. (**D**) Supplementing *ion* stimulation with bath-applied CabTRP Ia (10^-7^M) can rescue a crashed *C. magister* gastric mill rhythm at elevated temperature. Intracellular recording of LG during continuous extracellular *ion* stimulation with 13 Hz at 10 and 14°C. LG was rhythmically active at 10°C, but activity ceased at 14°C. Bath-applying CabTRP Ia re-established the rhythm at 14°C with unchanged *ion* stimulation.

We found similar results in *C. maenas*. As the temperature was increased, the LG resting membrane potential hyperpolarized (Fig. 3B). Concurrently, the amplitudes of its action potentials and the PSPs of the MCN1-like projection neuron diminished.

### Increasing descending projection activity or supplementing released neuropeptide rescues the gastric mill rhythm at high temperatures

We have previously shown that in *C. borealis*, MCN1 activity increases with higher temperature, and that increasing MCN1 firing frequency can rescue the gastric mill rhythm at high temperatures. We have also shown that high MCN1 firing frequencies correlate with ongoing gastric mill rhythms at high temperatures (Städele et al., 2015, DeMaegd and Stein, 2021, Städele and Stein, 2022). Also, when MCN1 activity is maintained at its cold activity level with a slow spike rate, supplementing the released neuropeptide with bath-applied CabTRP Ia selectively to the STG can also rescue the gastric mill rhythm at high temperatures. Figure 3C shows an original recording of the *C. magister* LG, with rhythmic bursting at 10 and 14°C after the rhythm crashed. Increasing *ion* stimulation frequency restored LG bursting and rescued the rhythm (N=7 of 7 tested animals). Similarly, when CabTRP Ia (10^-7^M) was bath-applied at 14°C instead of increasing *ion* stimulation frequency, the rhythm was also restored (Fig. 3D, N=7 of 7 tested animals). This suggests that the activity of descending projection neurons (and the release of neuropeptides from them) is also critical for temperature resilience of the gastric mill rhythm in this species. We noted that in no case was the application of CabTRP Ia alone sufficient to elicit a rhythm. Maintaining the *ion* stimulation was always necessary.

In *C. maenas* preparations where gastric mill rhythms could be elicited, higher *ion* stimulation frequencies also restored a crashed rhythm at high temperatures (Fig. 4A; N=3). Unfortunately, because of the small number of successful preparations, we could not test the effects of CabTRP Ia on the gastric mill rhythm in this species. However, we tested whether higher *ion* stimulation frequencies or CabTRP Ia application could elicit gastric mill rhythms in preparations where *ion* stimulation did not elicit a rhythm in the first place. This was not the case (N=8 for stimulation, N=3 for CabTRP Ia).

**Figure 4:**
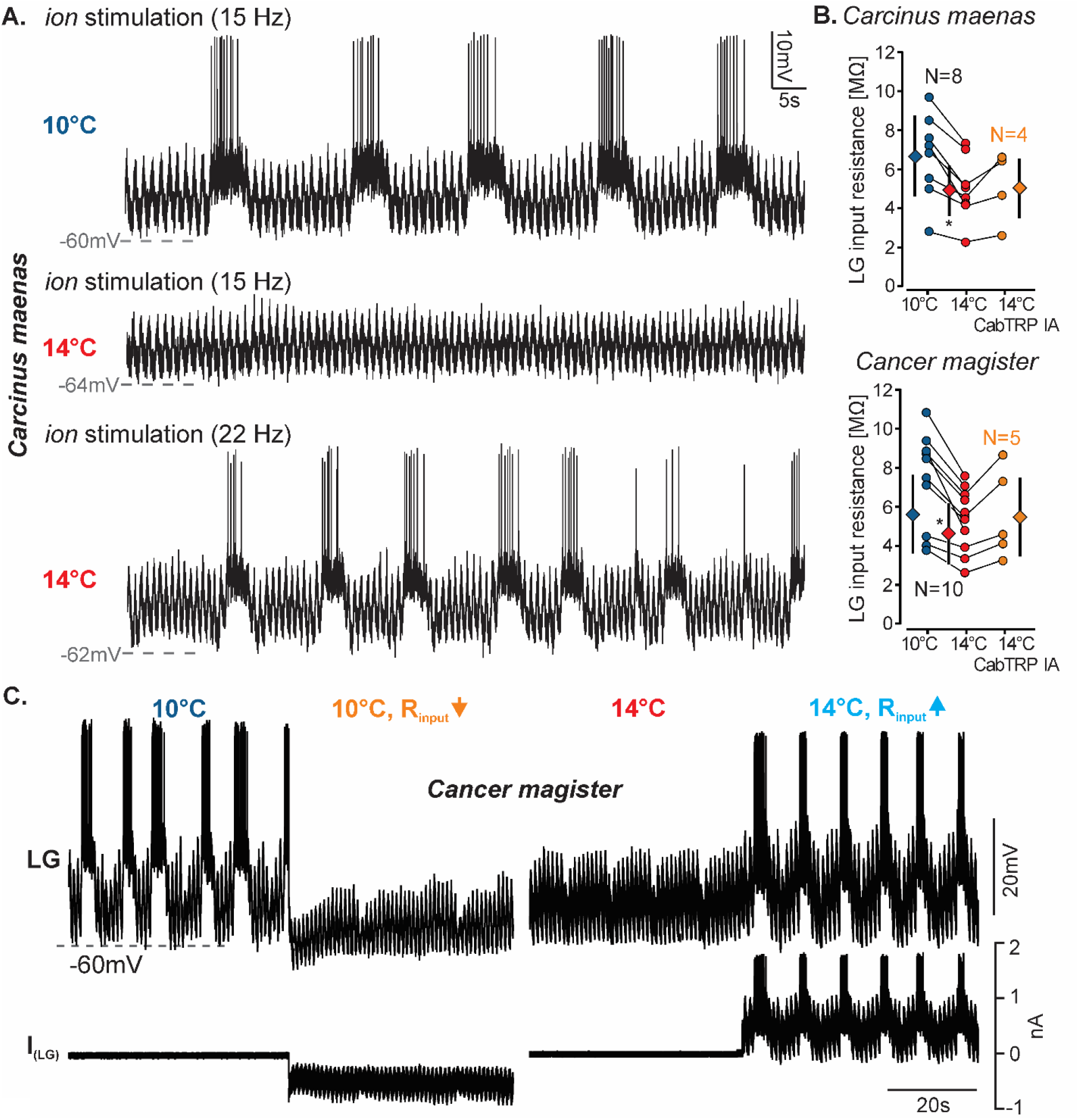
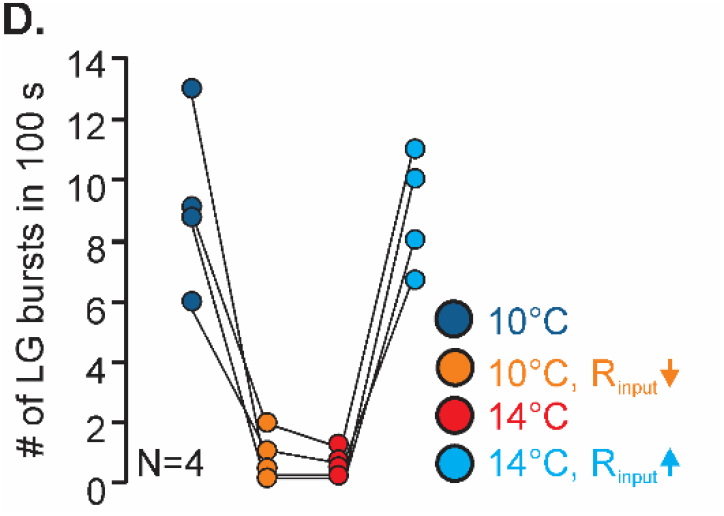
(**A**) Increasing *ion* stimulation frequency can rescue a crashed *C. maenas* gastric mill rhythm at elevated temperature. Intracellular recording of LG during continuous extracellular *ion* stimulation with 15 Hz at 10 and 14°C. LG was rhythmically active at 10°C, but activity ceased at 14°C. Increasing *ion* stimulation frequency to 22 Hz re-established the rhythm at 14°C. (**B**) Comparison of temperature effects on LG’s input resistance in all tested *C. magister* and *C. maenas*. Top: LG input resistance in *C. magister* at 10, 14, and 14°C with CabTRP Ia (10^-7^M) bath applied selectively to the STG. *P<0.05, One way repeated measures ANOVA, N=5; F(2,10)=5.018, Student-Newman-Keuls Method post-hoc test (P<0.05). Bottom: LG input resistance in *C. maenas:* *P<0.05, One way repeated measures ANOVA, N=8; F(2,10)=13.90, Student-Newman-Keuls Method post-hoc test (P<0.05). (**C**) Changes in input resistance are sufficient to terminate and rescue the rhythm in *C. magister* in 50% of the preparations tested: Intracellular recording of LG during continuous *ion* stimulation at 10°C. Rhythmic activity ceased when LG’s input resistance was artificially reduced by dynamic clamp (10°C +IR↓). At 14°C, rhythmic LG activity terminated even without artificial reduction of the input resistance. However, rhythmic activity was recovered when the input resistance was artificially increased (14°C+IR↑), rescuing the rhythm. (D) Quantification of LG bursts in the different dynamic clamp conditions.

### LG cell input resistance diminishes with increasing temperature

To understand the cellular cause of the gastric mill rhythm crash at high temperatures, we turned back to the cellular properties of LG. LG’s cell body and neuropil processes do not support regenerative action potentials (Raper, 1979). The diminishment of LG action potential and PSP amplitudes thus suggested that LG’s input resistance diminished with higher temperatures. To test this, we measured LG input resistance in two-electrode current clamp. In both species, input resistance was lower at higher temperatures (Fig. 4B). Because previous data from *C. borealis* had shown that CabTRP Ia modulation increases the neuronal length constant by counterbalancing low input resistance (DeMaegd and Stein, 2021), we measured the effect of CabTRP Ia on input resistance in a subset of the experiments (Fig. 4B, orange). CabTRP Ia (10^-7^M, selectively applied to the STG) restored input resistance at 14°C to values indistinguishable from saline values at 10°C in both crab species.

### Restoring input resistance rescues the gastric mill rhythm in *C. magister*

We tested whether the observed changes in input resistance are sufficient to explain the crash of the rhythm at high temperatures. For these experiments, we reduced input resistance at cold temperatures using dynamic clamp and measured whether this reduction stopped the rhythm. Conversely, we increased input resistance at high temperatures to test whether this increase was sufficient to maintain the rhythm at that temperature. These recordings were carried out in two-electrode dynamic clamp, with one electrode measuring voltage and the other injecting current. Changes in input resistance were matched to those measured during temperature changes in the same experiment (see Materials and Methods).

Figure 4C (top) shows an original recording of the gastric mill rhythm in *C. magister* in 4 conditions: at 10°C, at 10°C with dynamic clamp-reduced input resistance, at 14°C with no dynamic clamp manipulation, and at 14°C with dynamic clamp-increased input resistance. At 10°C, a regular gastric mill rhythm was observed. Increasing the temperature to 14°C terminated the rhythm. The decrease in input resistance observed during this temperature change was sufficient to terminate the gastric mill rhythm when applied through dynamic clamp at 10°C. Conversely, when input resistance was increased by the same amount at 14°C, the rhythm was restored. Figure 4D provides the quantitative data for four successful experiments. However, in contrast to *C. borealis*, changing input resistance by the amount suggested by the temperature increase was not always successful. In four other experiments (not shown), the rhythm at 10°C was not terminated by the change in input resistance. Conversely, the rhythm could also not be restored at 14°C. Instead, LG continued to produce bursts, albeit often with fewer action potentials and at lower frequencies. However, we noted that when we introduced larger changes in input resistance, the rhythm at 10°C could be terminated in all preparations, and restored at 14°C. Consequently, in these preparations, increasing the temperature was able to terminate the rhythm, but increasing the input resistance by the temperature-induced amount did not rescue the rhythm. This could suggest that additional cellular or synaptic mechanisms contributed to the heat-induced termination of the rhythm.

## Discussion

The main finding of our study is that, like in the previously tested *C. borealis*, the intrinsic permissive (‘stable’) temperature range of the gastric mill rhythm in *C. magister* and *C. maenas* was small (3-4°C). This was the case even though these species experience larger temperature fluctuations in their habitat (Fig. 1A), and their pyloric rhythm was maintained during all temperature challenges we applied. In *C. maenas*, for example, the gastric mill rhythm crashed more than 20°C below the pyloric rhythm (Stein et al., 2023). This appears to suggest that the animals cannot chew food at elevated temperatures, given that the gastric mill rhythm no longer functions. However, this does not seem to be the case. Adult *C. maenas*, for example, survive and presumably feed and digest well in habitats with temperatures well beyond the 14°C at which the gastric mill rhythm crashed at threshold stimulation in our experiments. Their primary habitat is the intertidal zone (Crothers, 1968), where they are exposed to the most extreme fluctuations in ocean temperature. While originating from the temperate waters of Europe (Klassen and Locke, 2007), they recently experienced a worldwide expansion and survive average temperatures ranging from 0°C to more than 35°C (reviewed by Young and Elliot, 2020).

The daily temperature variations they experience easily exceed the 4°C increase that was sufficient to crash the gastric mill rhythm. Accordingly, the temperature limits for feeding are estimated to be 5°C and 25°C (McGaw and Whiteley, 2012, McGaw and Curtis, 2013), with acclimation and selection pressures likely to play an essential role in achieving this broad range of feeding temperatures (Cuculescu et al., 1998). We also regularly observed feeding in our tanks at 20°C, even though the gastric mill rhythm failed at 14°C, and our dissections suggest that ingested food had been processed. Similar arguments can be made for *C. magister* and *C. borealis* (the original species in which the thermal protection of CabTRP Ia was found (Städele et al., 2015), although the temperature fluctuations these species experience are likely smaller than in *C. maenas*. They are found in benthic, subtidal, and intertidal zones and show strong acclimation and thermotactic behaviors (Prentice and Schneider, 1979, Lewis and Ayers, 2014).

In *C. borealis, in vivo* recordings have shown that gastric mill rhythms can occur at temperatures higher than the crash temperature during threshold *ion* stimulation (Städele et al., 2015). Our study shows a possible solution for the apparent contradiction between small intrinsic and considerable behavioral temperature robustness: The gastric mill rhythm was maintained at more elevated temperatures when extrinsic modulatory activity was increased, or additional neuropeptide was bath-applied. This suggests that temperature robustness in the gastric mill circuit is highly dependent on extrinsic modulatory input in these species. However, temperature robustness also requires increased neuromodulator release when temperature rises. There is evidence for such an effect in *C. borealis*: MCN1 is the descending projection neuron that releases CabTRP Ia (Christie et al., 1997), and its firing frequency has been shown to increase with temperature in at least two conditions: spontaneous activity (Städele et al., 2015) and sensory-induced during gastric mill rhythms (Städele and Stein, 2022). Although the definitive test to determine if higher temperatures result in increased CabTRP Ia release is still pending, these studies strongly suggest this.

Model predictions have suggested potential mechanisms that explain the difference in temperature robustness between pyloric and gastric mill rhythms: The pyloric central pattern generator relies upon the activity of a continuously active pacemaker and strong inhibitory feedback within the pyloric circuit (reviewed by Stein, 2017). Activity in pyloric follower neurons is facilitated by a rebound from inhibition, and inhibitory feedback from follower neurons to the pacemaker contributes to pyloric cycle frequency control (Thirumalai et al., 2006). Weakened inhibition makes it susceptible to failure (Selverston et al., 2000). The pyloric circuit thus shows features of a so-called escape circuit. Modeling studies predict that escape circuits possess a higher intrinsic robustness to temperature perturbations and rely less on extrinsic modulation (Morozova et al., 2022). In contrast, in the gastric mill circuit, LG cannot escape the Int1 inhibition but is instead released (disinhibited) when Int1 is inhibited by the pyloric pacemakers (Marder et al., 1998, Bartos et al., 1999). Release circuits are predicted to be more intrinsically sensitive to rising temperatures, but also to continue to function at these high temperatures when extrinsic neuromodulation is available. In that modeling study, the ionic current evoked by CabTRP Ia (I_MI_) was tested, and its previously demonstrated effect of opposing leak currents and amplifying oscillations in feedback circuits (either synaptic or cell-intrinsic) was shown to be a likely candidate for enabling temperature robustness (Morozova et al., 2022).

A consistent effect of elevated temperature in the two species we tested was the hyperpolarization of the resting membrane potential and a decrease in input resistance. The latter led to a shunt of postsynaptic potentials and action potentials, as evidenced by their diminished amplitude at elevated temperatures. Temperature effects on input resistance are well described in other systems as well. For example, the input resistance in goldfish Mauthner cells is consistently smaller after acclimation to elevated temperatures (Szabo et al., 2008). Changes in input resistance are known to play an essential role in network oscillations (Cymbalyuk et al., 2002, Blethyn et al., 2006, Koizumi et al., 2008, Zhao et al., 2010), regulation of excitability (Rekling et al., 2000, Brickley et al., 2007) and switches in activity states of neurons (Gramoll et al., 1994). Our results suggest that in all tested crab species so far, including *C. borealis* (DeMaegd and Stein, 2021), input resistance contributes to the termination of the rhythm at elevated temperatures. However, our dynamic clamp experiments also suggest that additional factors may add to the termination of the rhythm in *C. magister* because changing the input resistance was insufficient to recreate the temperature effects in all experiments. Additional factors may include effects on synaptic properties, voltage-gated ionic currents, and other neurons that are part of the gastric mill central pattern generator. The detailed connectivity of the gastric mill circuit in *C. maenas* and *C. magister* is unknown. However, in both species, LG’s membrane potential trajectory during the gastric mill rhythm and its synaptic inputs look very similar to the well-characterized gastric mill circuit of *C. borealis* (Coleman et al., 1995, Stein et al., 2007). Yet, we observed a few differences. Most notably, in *C. magister*, there were slow periodic subthreshold hyperpolarizations during failed rhythms (Fig. 3C) that were absent in *C. maenas* (Fig. 4A) and *C. borealis* (Coleman et al., 1995, Stein et al., 2007).

The identity of the descending projection neuron that activates LG during *ion* stimulation has also not been confirmed in *C. maenas* and *C. magister*. It is known that synaptic connectivity between homolog neurons can vary between species, as can the identity and actions of peptide co-transmitters (summarized in (Stein et al., 2016). Surprisingly, however, increasing the descending projection neuron’s activity through *ion* stimulation restored the rhythm in all preparations. This indicates that whatever differences exist between species and contribute to LG’s temperature crash can be counterbalanced by descending modulation. Strikingly, the application of MCN1’s peptide co-transmitter CabTRP Ia was able to rescue the *C. magister* gastric mill rhythm at elevated temperature, and we suspect that *in vivo*, the additional peptide is provided by higher projection neuron activity. A remote possibility is that temperature effects on the ionic current elicited by CabTRP Ia (I_MI_) could also contribute to temperature robustness. Its conductance may increase with temperature, boosting temperature robustness by *counterbalancing* negative leak conductance (Zhao et al., 2010), thus restoring gastric mill oscillations. However, how much I_MI_ changes with temperature remains to be determined. Surprisingly, our dynamic clamp experiments succeeded in only half of the experiments in *C. magister*. In the other half, the injected conductance was insufficient to mimic the temperature effects, and the gastric mill rhythm continued, albeit with weaker bursts. This may reflect space-clamp problems in our approach. LG in *C. magister* is a very large cell. We estimate its cell body to be >100µm, with a primary neurite of at least the same length. This could interfere with the ability of our (somatic) clamp to reach the distant dendritic processes of the cell where synaptic input arrives and the spike initiation site where action potentials are generated. This may be particularly the case at elevated temperatures because input resistance drops (Fig. 4B), suggesting that the length constant of the neuron shortens. This effect was present in LG of *C. borealis* (DeMaegd and Stein, 2021).

Counterbalancing detrimental temperature effects on the nervous system is critical for maintaining neuronal function and is often essential for survival. This is particularly true when circuits that drive vital body functions are affected. Central pattern-generating circuits are ubiquitously found in the nervous systems of all animals. It is striking how much even small temperature changes can alter the output of these vitally essential circuits. Hyperthermia during fever or heat stroke can, for example, cause failures of the respiratory pattern generator and induce apnea (Tryba and Ramirez, 2004). Generally, it is assumed that the permissible temperature range a neuronal circuit displays is related to the temperature fluctuation it experiences (Stein and Harzsch, 2021). For homeotherms, including mammals, the permissive range is predicted to be small, given the relatively small variations in body temperature that occur daily or monthly. In contrast, the permissive range is expected to be much more significant for poikilotherms, particularly those that live in habitats with large temperature fluctuations. Recent studies have provided some evidence for this, showing that the pyloric rhythm of intertidal species possesses a more extensive permissive range than in deeper-ocean species (Soofi et al., 2014, Stein et al., 2023). This suggests that neuronal mechanisms to counterbalance temperature effects are more sophisticated in animals that experience large temperature fluctuations in their environment. It also indicates that neuronal activity in homeotherms may be more susceptible to temperature changes. This becomes particularly important when neuronal circuits are studied at temperatures that are very different from the actual internal temperature of the animal, as is common practice in many fields. For example, the pyloric and gastric mill central pattern generators have been studied for over six decades, and they have provided much insight into the functioning principles of neuronal pattern generation and its modulation. However, most studies were conducted at an agreed-upon constant temperature that did not account for the temperatures the nervous system is exposed to *in vivo*. As a consequence, the finding that the intrinsic temperature robustness of the pyloric rhythm is large (Tang et al., 2010) and that of the gastric mill rhythm is small (Städele et al., 2015) was initially controversial. It seemed contradictory that two parts of the same neuronal system (the stomatogastric nervous system) would respond differently to temperature changes. Our findings resolve this contradiction: the lack of intrinsic robustness of the gastric mill rhythm is rectified by the effects of extrinsic modulation, which largely enhances temperature robustness. The pyloric and gastric mill central pattern generators thus serve as an extreme example of how fundamentally different neuronal circuits can respond to temperature changes. They are also a reminder that dramatic shifts in circuit activity may occur when circuits are exposed to changing temperatures and modulatory conditions.

## Acknowledgments

We would like to thank Dr. Markus Frederich (University of New England, Maine, USA) and Adam Smith (Perfect Limit Green Crabs, Massachusetts, USA) for the *Carcinus maenas* used in this study. We would also like to thank Dr. Margaret DeMaegd (New York University) for providing some of her input resistance data.

## Data availability

All relevant data can be found within the article.

